# Non-specificity fingerprints for clinical stage antibodies in solution

**DOI:** 10.1101/2023.02.13.528263

**Authors:** Therese W. Herling, Gaetano Invernizzi, Hannes Ausserwöger, Jais Rose Bjelke, Thomas Egebjerg, Søren Lund, Nikolai Lorenzen, Tuomas P. J. Knowles

## Abstract

Monoclonal antibodies (mAbs) have successfully been developed for the treatment of a wide range of diseases. The clinical success of mAbs, does not solely rely on optimal potency and safety, but also require good biophysical properties to ensure high developability potential. In particular, non-specific interactions are a key developability measure to monitor during discovery. Despite an increased focus on the detection of non-specific interactions, their physicochemical origins remain poorly understood. Here, we employ solution-based microfluidic technologies to characterise a set of clinical stage mAbs and their interactions with commonly used non-specificity ligands to generate non-specificity fingerprints, providing quantitative data on the underlying physical chemistry. Furthermore, the solution-based analysis enables us to evaluate the contribution of avidity in non-specific binding by mAbs. Based on our findings, we propose a quantitative solution-based non-specificity score, which can be exploited in the development of biological therapeutics and more widely in protein engineering.

---

Monoclonal antibodies are prominent amongst the current best-selling pharmaceuticals. These versatile biological drugs have been optimised to target a wide range of conditions including autoimmune, cancer, and infectious diseases. Features such as high target affinity and specificity, biocompatibility, immune effector functions and desirable pharmacokinetic properties are factors leading to antibodies becoming one of the preferred therapeutic modalities. However, the development process is costly and often unsuccessful in delivering a commercial product.^1,2^ In spite of their vast potential, the 100^th^ mAb was only approved recently.^3^ Antibody hits identified during discovery are often subjected to affinity maturation campaigns using display-based platforms. Recent studies have demonstrated that these maturation campaigns can compromise mAb development through a trade-off between affinity and specificity.^4–7^ The propensity for non-specific interactions emerges as a critical determinant of developability i.e. the suitability of antibody drug candidates to succeed as medicines, Figure 1.^5,8,9^ Consequently, screens for non-specificity are rapidly becoming industry standard and are implemented in parallel with functionality measures during the discovery and optimisation phases.^5,8–12,23^

**Figure 1:**
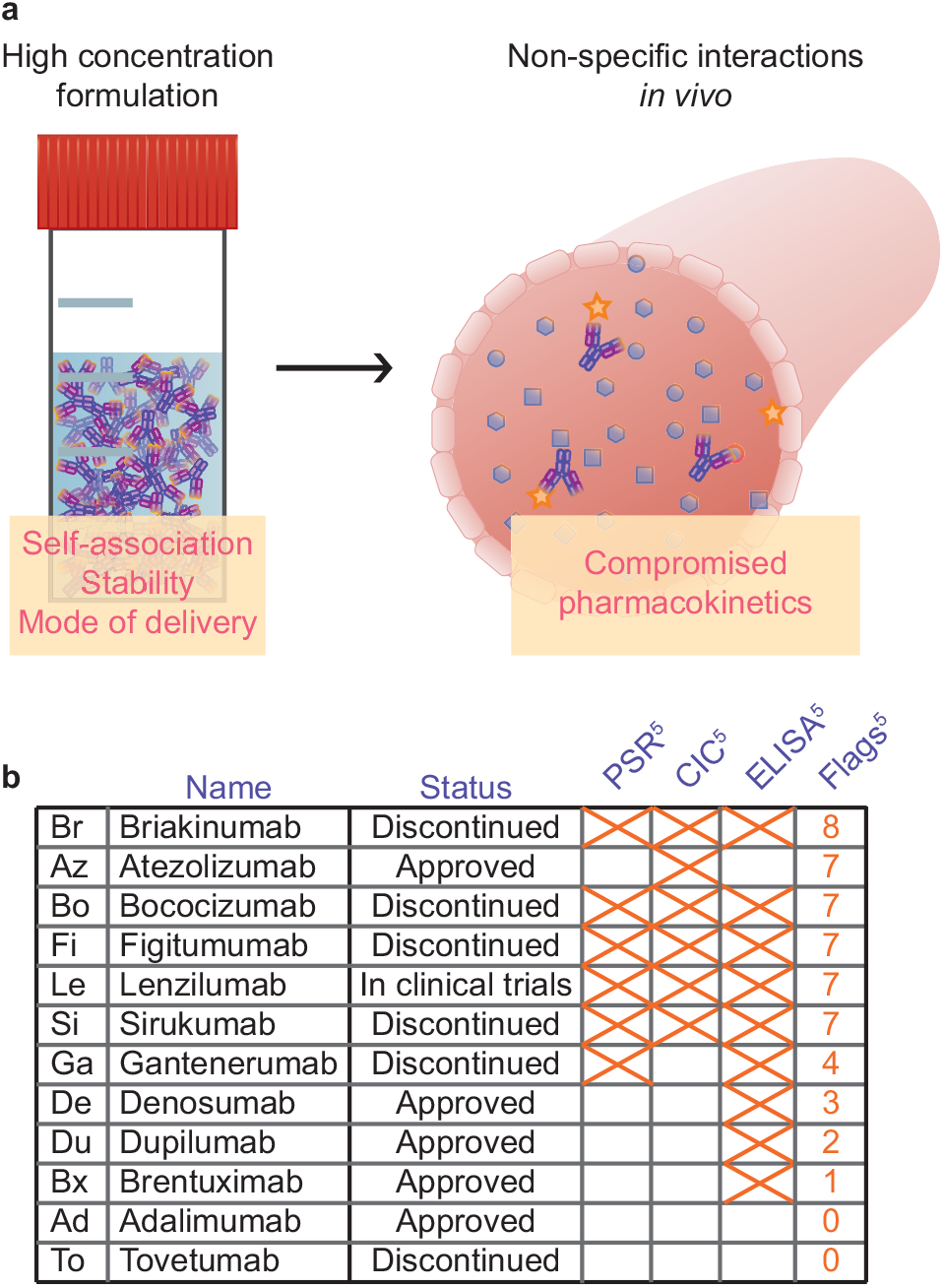
**a** Left, high mAb concentration in low complexity solutions. Right, low mAb concentration under crowded conditions in a complex solution. **b** Panel of clinical stage antibodies ranked according to their reported biophysical properties. Flags detected in specific assays in reference 5 are indicated with an X for poly-specificity reagent (PSR), cross-interaction chromatography (CIC), and enzyme-linked immunosorbent assay (ELISA) The number of flags out of 10 is displayed.^5^

Remarkably, in spite of the current focus on screening for non-specificity as a key developability parameter, the underlying physical chemistry remains largely unexplored.^5,8,13^ Direct measurements of non-specific binding events in solution have not been within the scope of mAb development campaigns, and traditional assays are poorly suited to address these questions, as they rely on indirect reporters, such as light scattering, and potentially disruptive processes such as surface attachment or for samples to pass through a matrix.^5,22,23^ Here, we address the need for quantitative solution-based non-specificity assays by applying microfluidic techniques and we generate non-specificity fingerprints for a panel of clinical stage antibodies by challenging them with a range of physiologically relevant ligands, Figure 1 and SI Figure I.^14–16^ Crucially, by evaluating changes to physicochemical parameters, such as the size and charge of antibodies, we detect interactions in a quantitative manner without surface attachment.^15–19^

The bivalent antibody format permits ligand binding avidity, which could exacerbate nonspecific interactions in surface-based assays against an immobilised ligand (CIC, ELISA and similar).^22,23^ The solution-based platform provides us with the opportunity to gain a better understanding of the role of avidity in determining mAb affinity and how this effect may translate to *in vivo* situations e.g. as a driver of binding to interfaces such as the endothelial cell surface.^23^ We find that avidity is a key promoter of non-specific binding, and we evaluate the contribution of avidity potential to the apparent *K_d_* by exposing a mAb to DNA oligomers of decreasing length.

We propose that non-specificity issues can be identified through changes to the ligand size or charge, this approach yields a score, which can be evaluated as a stand-alone measurement without the need for a cohort dataset as benchmark. Taken together, we establish microfluidics as a platform for the quantitative, in-solution characterisation of biologics. Furthermore, we provide crucial insights into the impact of avidity on non-specific binding and rationalise the physicochemical drivers that promote these interactions, enabling these features to be addressed during the development process.

## Results and discussion

### Physicochemical properties of monoclonal antibodies

A dozen mAbs are selected based on their reported biophysical properties, Figure 1**b** and SI Table 1.^5^ In particular, we have chosen those with a high number of developability flags in past studies (Briakinumab, Atezolizumab, Bococizumab, Figitumumab, Lenzilumab and Sirukumab). Brentuximab, Dupilumab and Denosumab have mainly been flagged in surface-based non-specificity screens, and we were interested in discovering whether these non-specific interactions translated to solution-based assays. Additionally, we have included Adalimumab and Tovetumab as references with good reported biophysical properties, i.e. we would expect a low propensity for non-specific binding in these antibodies.^5^ In addition, the mAb selection is designed to include sequence charges ranging from the near-neutral Brentuximab to the highly positively charged Lenzilumab.^5^

Microfluidic technology enables us to interrogate key solution properties of mAbs, and we screen the diffusion coefficient (*D*), the corresponding hydrodynamic radius (*R_H_*), the electrophoretic mobility (*μ_e_*), and the effective solution charge (*z_e_*), SI Table 1.^14–17,19^ Notably, we screen unlabelled antibodies, SI Figure 1. This feature allows us to not only measure the nonspecific interactions, but also to perform quality control on our antibody and ligand stocks.

Antibody self-association is a common issue that can potentially compromise the reliability of interaction studies, measurements of the mAb *R_H_* can detect this process.^11,20,21^ We screen for non-specific interactions at a relatively low mAb concentration (1 mg/ml) in order to limit the risk of mAb self-association. No significant increase in mAb size was observed under native solution conditions at physiological salt levels (hs, 150 mM NaCl, 20 mM Hepes-NaOH pH 7.4) or low salt (ls, 15 mM NaCl, 2 mM Hepes-NaOH pH 7.4), SI Table 1. All samples and buffer solutions contain 0.01% v/v Tween20 to prevent surface adhesion within the microfluidic channels.

### Avidity enhances non-specific interactions

Screens for non-specific interactions are an integral part of the drug development pipeline.^5,8–12^ However, commonly used non-specificity assays require surface attachment or for the samples to pass through a matrix, conditions that favour avidity.^22,23^ The ability to tune the avidity potential e.g. to reflect membrane association or interactions with soluble ligands is key for developing an accurate understanding of the non-specificity profile for mAbs.

When administered as drugs, mAbs encounter insulin and cell-free DNA fragments in circulation.^24–26^ A propensity for non-specific interactions with either biomolecule could thus affect the function of mAbs through competition from off-target interactions.^5,7–12^ Circulating cell free DNA can be double-stranded or single stranded.^25^ DNA and insulin are two frequently employed non-specificity probes, and antibodies that interact strongly with these negatively charged biomolecules have been reported to have an increased risk of poor in vivo half-lives.^10^ Therapeutic mAbs thus need to achieve a balance between typically positive sequence charges to facilitate interactions when targeting membrane proteins and avoidance of non-specific interactions e.g. due to charge patches on the mAb surface.^9^

We use microfluidic diffusional sizing to record the *R_H_* for mixtures of mAbs and monomeric insulin-CF488 (1 *μ*M, icf) or single-stranded DNA oligomers labelled with Cy3 (20mer, 1 *μ*M) at physiological salt levels, Figure 2**a-b** and SI Figure 2. Fluorophore-labelling of the non-specificity ligands enables the size of both the unlabelled mAbs and the targets to be measured in each sample. Unlike surface-based assays, the analyte concentrations in solution are well-defined and can be tuned to investigate specific binding affinities. Microfluidic technologies are compatible with high-viscosity solutions, so non-specificity assays could also be conducted at elevated protein concentrations, in serum, or cell lysate.^15^

**Figure 2:**
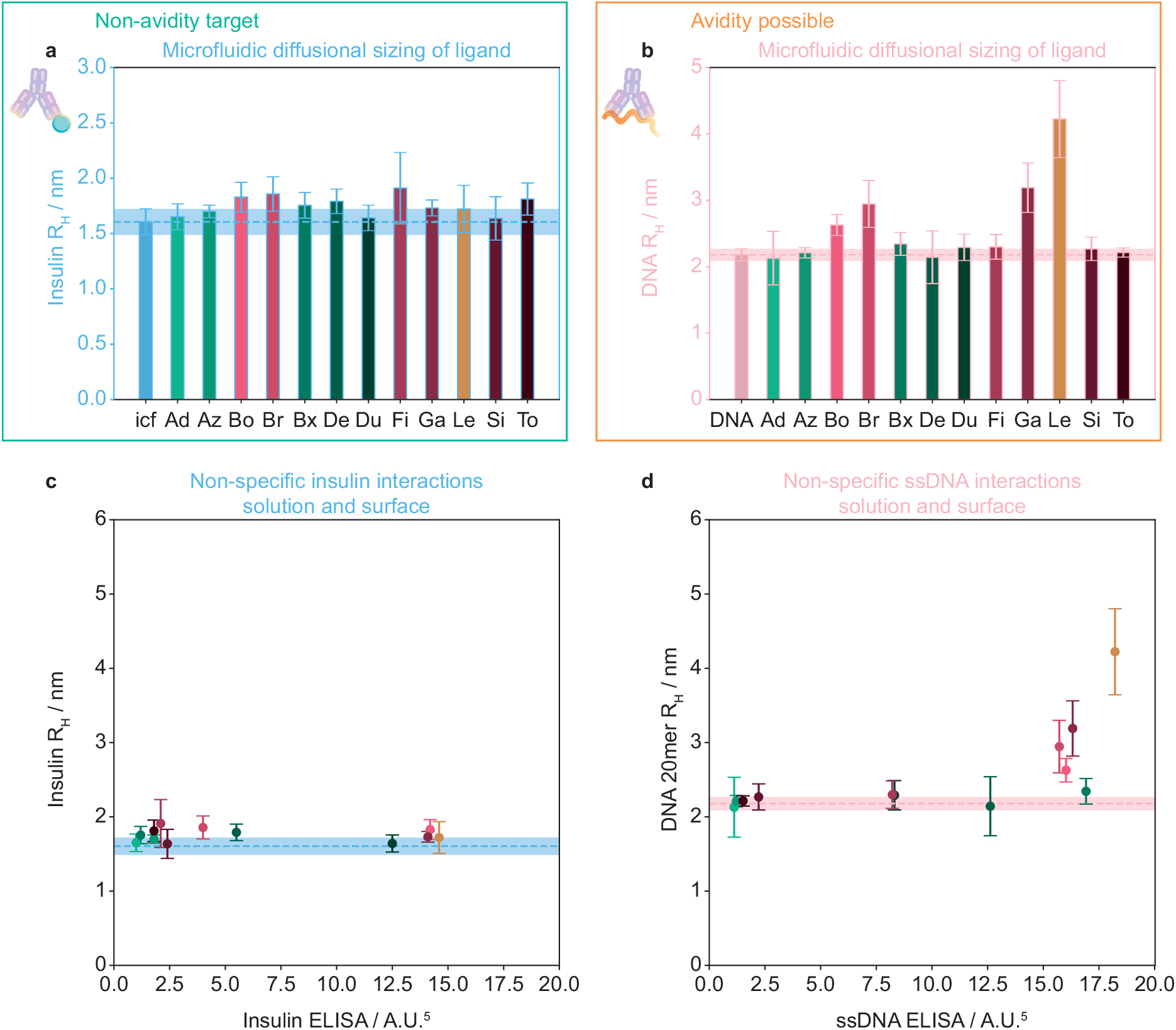
**a** Non-specific binding assayed in free solution under non-avidity conditions in the high salt buffer between unlabelled mAb (1 mg/ml, 6.7 *μ*M) and 1 *μ*M monomeric insulin-CF488 (icf, blue, error bars show ± standard deviation for three independent repeats). **b** Interactions where target avidity is possible with a single-stranded DNA polymer (20mer, 1 *μ*M, pink, Cy3-labelled). **c** The observed *R_H_* for 1 *μ*M insulin against the reported ELISA signal for each mAb against immobilised insulin (colours as in **a-b**).^5^ **d** The observed DNA *R_H_* against the reported ELISA signal for each mAb against ssDNA (colours as in **a-b**).^5^

No significant interaction between the antibodies and insulin was observed in solution, Figure 2**a**. ELISA screens were performed against immobilised ligands, and avidity effects would result in low effective dissociation rates, thus increasing the apparent interaction affinity. Negatively charged polymers such as DNA provide an opportunity to investigate non-specific binding under avidity conditions, SI Figure 2 and Figure 2**b+d**. In the surface-based assay most of the mAbs appear to bind insulin and DNA to some extent. However, the solution-based screen only flags a subset of the antibodies, Figures 1**b** and 2**c-d**. In particular, the observed DNA *R_H_* increases in the presence of Bococizumab, Briakinumab, Gantenerumab, and Lenzilumab indicating binding. This result is in good agreement with ELISA data by Jain et al.^5^ where these four mAbs are amongst the five with the highest ELISA signal against ssDNA (Figure 2**d**). Notably, none of the mAbs flagged in the solution-based measurements have received regulatory approval yet. In a further validation of our approach, we compare our data with the results reported by Kraft et al. for their column-based heparin interaction assay, SI Figure 3.^23^ We find that they measured the highest relative heparin retention times for the same four mAbs, which we highlight here as prone to non-specific interactions.

Three of the approved mAbs, Brentuximab, Denosumab, and Dupilumab, were selected for this study because they were flagged as poorly behaved in surface-based assays (ELISA and baculovirus particle), Figure 1**b**.^5^ We were interested in whether the reported interactions would also be observed without surface-attachment. In solution, neither ligand showed an *R_H_* increase in high salt buffer with these mAbs, Figure 2**a-b** and SI Figure 2.

Lenzilumab led to the largest DNA size increase, Figure 2**b** and SI Figure 2**b**. We therefore set out to determine the role of binding avidity in promoting this interaction, Figure 3. To this end, we measured the *R_H_* of 100 nM Cy3-labelled DNA oligomers (20mer, 10mer and 6mer) as a function of Lenzilumab concentration. Condensation to form micron-scale particles was observed for a DNA 100mer, SI Figure 4**b**. Here, we consider the interactions as Fab-DNA binding, as this fragment is varied between the antibodies in our screen, whereas they share an IgG1 Fc. Le had micromolar affinity for the 20mer (*K_d app_* = 3.4 *μ*M per antigen binding site - two per mAb), and clusters were formed (*R_complex_* = 10.7 nm > *R_Le_*). The concentration of DNA used here is comparable to those of cell-free dsDNA in plasma (1-10 ng/ml in healthy individuals, rising to 100s of ng/ml in cancer patients),^26,27^ and a significant fraction is bound at a typical mAb *c_max_* of 0.1 mg/ml (0.67 *μ*M) in plasma.^28^ This type of non-specific interaction could thus be highly significant for the function of therapeutic antibodies.

**Figure 3:**
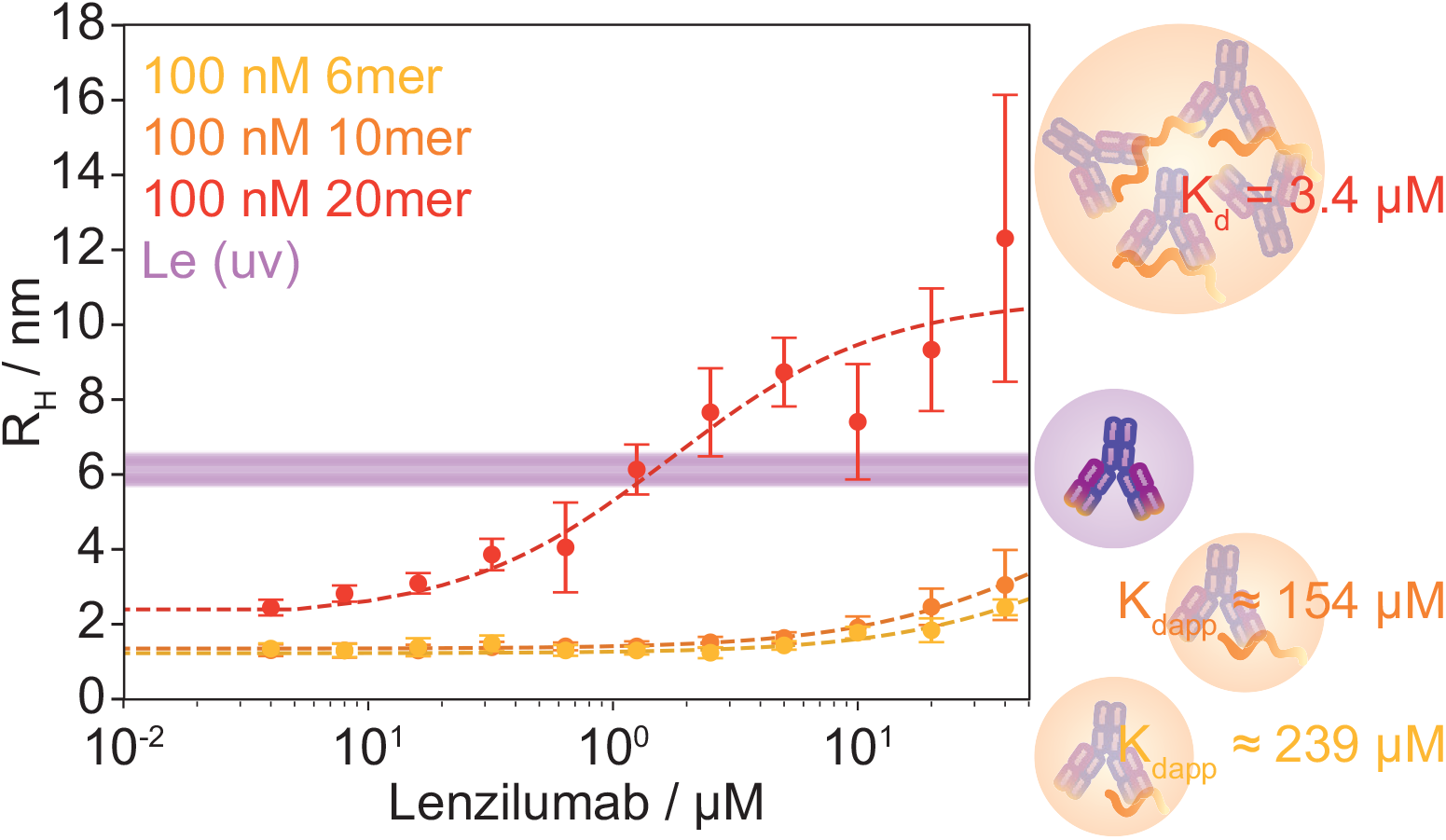
Diffusional sizing of 100 nM Cy3-labelled DNA oligomers as a function of the concentration of the mAb Lenzilumab at physiological salt concentrations. Three oligomer lengths 20 (20mer, red), ten (10mer, orange) and six bases (6mer, yellow) were investigated to evaluate the polymer size requirement for avidity effects in non-specific mAb-DNA interactions. Even the relatively short 20mer formed higher order complexes (*R_complex_* = 10.7 nm > *R_mAb_*) with a *K_d app_* of 3.4 *μ*M per antigen binding site (assuming two per antibody). When binding avidity was prevented by reducing the oligomer length to ten or six bases, we observed a dramatic decrease in binding affinity, by an estimated two orders of magnitude (*K_d app_* ≈ 154 and 239 *μ*M respectively). The antibody *R_H_* was measured for all sample compositions with Le ≥ 640 nM and found to be constant within error (purple shaded areas show average ± standard deviation for the three datasets). Right, schematic illustrations and *K_d app_*.

For mAb concentrations of 640 nM (≈0.1 mg/ml) and above, the antibody in the sample mixture was sized via intrinsic protein fluorescence. The antibody is in excess at the plateau values so the complexes are likely to involve multiple mAbs cross-linked by the DNA oligomer. The constant *R_mAb_* in these samples indicates that DNA binding does not induce aggregation of the whole Lenzilumab population.

Remarkably, when the polymers were shortened to prevent ligand avidity, the apparent Lenzilumab affinity was reduced by approximately two orders of magnitude (*K_dapp_* ≈ 154 and 239 *μ*M respectively when *R_complex_* was fixed to *R_mAb_*). Similarly to monomeric insulin, charge complementarity alone is not sufficient for high affinity non-specific interactions at physiological salt concentrations. Avidity therefore emerges as a key factor promoting off-target mAb interactions.

### Screening for non-specific interactions

The microfluidic platform for quantitative biophysical characterisation of mAbs provides a versatile measure of non-specific interactions under a wide range of solution conditions. We create a non-specificity fingerprint for the mAbs by monitoring the change in *R_H_* for mAb-ligand mixtures relative to the individual components at high salt (150 mM NaCl) and low salt (15 mM NaCl), Figure 4**a-b**. We challenge the mAbs with molecules that may be encountered *in vivo*, including negatively charged polymers (1 *μ*M ssDNA oligomers and 1 mg/ml heparin), a small negatively charged protein (1 *μ*M monomeric insulin), and human serum albumin (2 mg/ml HSA), which carries both a negative charge and has exposed hydrophobic patches.^29,30^ At 35-50 mg/ml, HSA is the most abundant protein in plasma, and it interacts with a wide range of drug molecules. Indeed, binding to HSA is used as a strategy to extend the half-life of medicines.^30^ Only HSA has significant levels of intrinsic fluorescence at the concentrations used here, the observed *R_H_* is therefore a weighted average of the mAb and HSA contributions.

**Figure 4:**
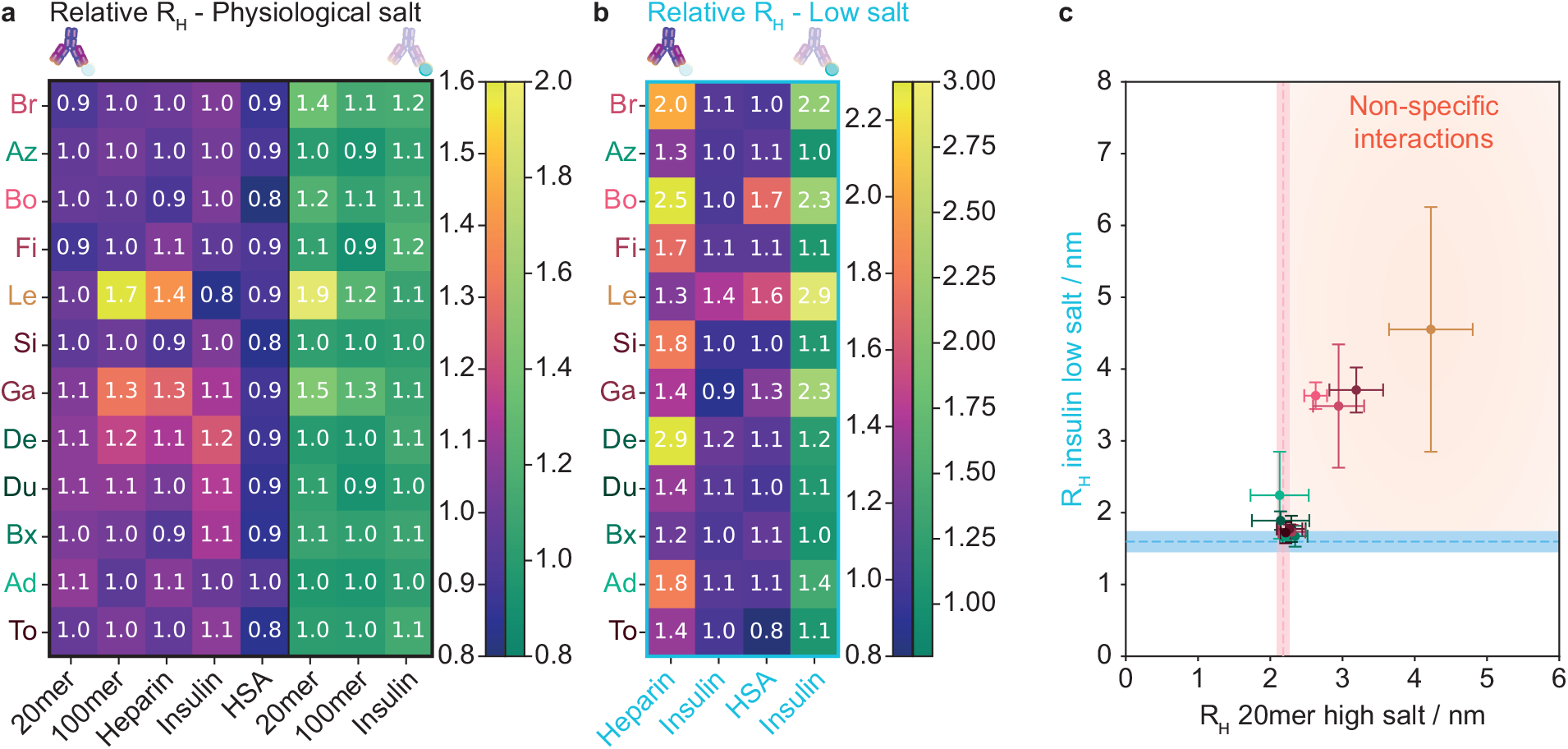
**a** The relative size of the antibodies (purple) in the presence of non-specificity probes in high salt buffer. Green data is relative *R_H_* for labelled non-specificity ligands. An average size is recorded for HSA and mAb samples using intrinsic fluorescence. Antibodies are ordered according to number of reported flags^5^ as in Figure 1**b**, and labels are colour coded by approval status (green = approved, brown = in clinical trials, pink = discontinued). **b** Relative size change for mAbs (purple) and non-specificity ligands (green) in low salt buffer. **c** A plot of *R_H_* for a ligand with avidity at high salt (ssDNA 20mer) against a non-avidity ligand at low salt (monomeric insulin). Four mAbs emerge as non-specific binders against both ligands (Bo, Br, Ga, Le).

Based on their advanced development stage, most of the mAbs investigated here would not be expected to show non-specificity at high salt, Figure 4**a**. Electrostatic interactions are promoted in the low salt buffer, and we observe extensive interactions with the negative polymers, e.g. with heparin, Figure 4**b**). For DNA, precipitation is observed upon dilution of the buffer (SI Figures 4 and 5), we therefore only probed for interactions with DNA at physiological salt levels.

Denosumab is marketed as an IgG2 rather than IgG1 antibody.^31^ The different isotype used in this study may be less stable than the antibody drug, resulting in poorer biophysical properties and non-specificity behaviour. This antibody is flagged for interactions with non-specificity ligands in the ls buffer and has a slightly increased size in the intrinsic fluorescence measurements, Figure 4.

We note that four of the mAbs were consistently flagged as prone to non-specific interactions (Bo, Br, Ga, and Le), Figure 4**c**. Notably, none of the approved mAbs were included in this group, these would have been considered false positives in our assay SI Figure 6. In order to identify accurately mAbs with a propensity for non-specific interactions, we therefore propose the use of the average relative *R_H_* for two or more ligands as a measure of non-specificity, SI Figure 6**b**.

#### Charge property evaluation

The charge properties of mAbs are important for target recognition, potency, pharmacokinetics, formulation, and as a quality control measure e.g. for post-translational modifications.^32,33^ Current strategies for assessing these use column chromatography to fractionate samples followed by mass spectrometry or capillary electrophoresis to assess the sample charge.^32,33^ Mass spectrometry in particular has limited buffer system compatibility, and column separation techniques may require non-physiological conditions, such as low pH, to operate optimally.^33^ The microfluidic platform employed here is compatible with common biological buffer systems and can be applied to determine *z_e_* under native solution conditions, Figure 5**a**.^17^ While the free-flow electrophoresis step is preferentially performed in low ionic strength buffers for optimum separation, microfluidics is a highly modular technology, and a desalting element can be included upstream of the electrophoresis channel.^34^

**Figure 5:**
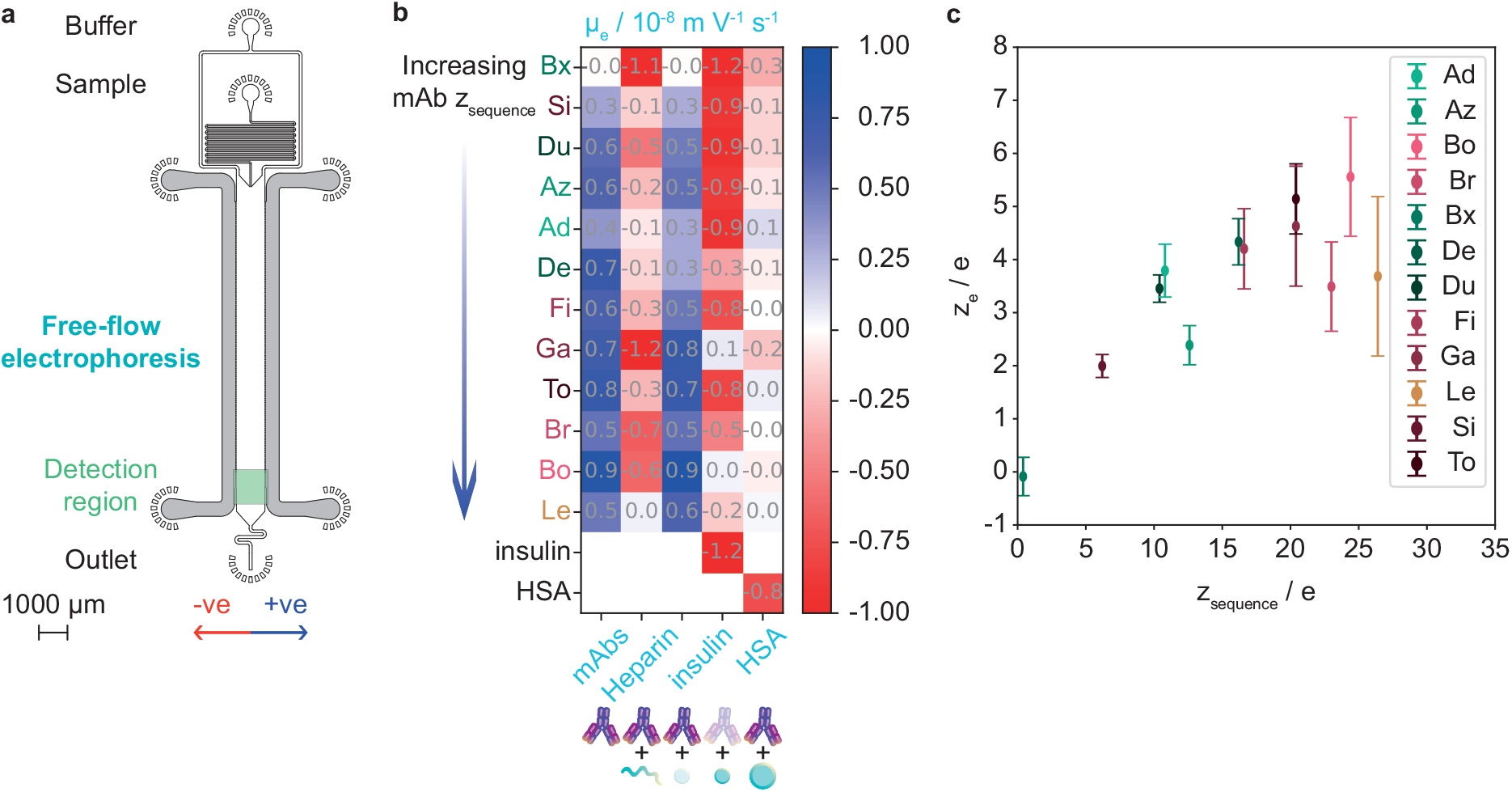
**a** Microfluidic free-flow electrophoresis design. Solid electrodes (grey) are incorporated along the main channel as previously described.^14^ An electric field is applied perpendicularly to the direction of flow, and sample molecules migrate according to their electrophoretic mobility. **b** Heat map of the electrophoretic mobility for the mAbs (1 mg/ml) in isolation and in the presence of non-specifcity targets heparin (1 mg/ml), monomeric insulin-CF488 (icf, 1 *μ*M), and human serum albumin (HSA, 2 mg/ml). **c** Effective charge (*z_e_*, ls) versus the dry sequence charge for the panel of mAbs investigated in this study.^5^

We have selected the antibodies in this study to have a wide distribution of sequence charges, from the near-neutral Brentuximab to the highly positively charged Lenzilumab.^5^ The *μ_e_* reports on the sample size to charge ratio and is therefore a useful parameter for detecting interactions where the complex *M_w_* is similar to that of the observed protein, for instance mAb versus mAb (150 kDa) +heparin (8-25 kDa). We measure *μ_e_* for individual mAbs and mixtures with nonspecificity ligands in low salt buffer, SI Table 1 and Figure 5**b**.

The mAbs bind insulin in low salt buffer, and it is therefore a useful non-specificity probe (Figures 4**b-c** and 5**b**). Most insulin interactions have little effect on the overall mAb *R_H_* and *μ_e_*. While there is little interaction with HSA in hs buffer, both electrophoresis and diffusional sizing show binding between HSA and the majority of mAbs under ls conditions, Figures 4**b** and 5**b**. The mAbs interact extensively with heparin at low ionic strengths. The change in *M_w_* would be small for mAb binding to a single polymer, however **R_H_** measurements reveal that clusters are formed, Figures 4**b** and 5**b**. The increase in charge complementary interactions at low ionic strength, indicates that these are driven by electrostatics.

We calculate *z_e_* based on *R_H_* and *μ_e_* and find that while *z_e_* increases with sequence charge, the absolute value is lower, Figure 5**c**.^17^ Charge-screening through interactions with solvent molecules and within the protein structure can contribute to the lower solution charge.^35^ Additionally, charged residues may be buried within the immunoglobulin folds, form salt bridges, or have suppressed pK_A_ values, e.g. due to clustering of charged residues. The antibodies are all expressed on the same IgG1 scaffold, so differences in overall charge originate mainly in the variable regions and choice of *κ* or λ light chains.

Four of the five mAbs with the highest *z_e_* and sequence charges are highlighted as prone to non-specific interactions, Figures 4**c** and 5**c**. High levels of positive charge have been associated with faster clearance rates *in vivo*, there is thus a trade-off between antibody potency and pharmacokinetics.^36^ It may therefore be desirable to avoid or modulate excessive positive charges during the development process (Figure 5**c** and 6). Notably, antibodies with low (Brentuximab) and moderately high (Denosumab, Dupilumab) *z_e_* have received approval as drugs. This observation indicates that the charge distribution in the antibody structure can play a key role in mAb developability.^9^

**Figure 6:**
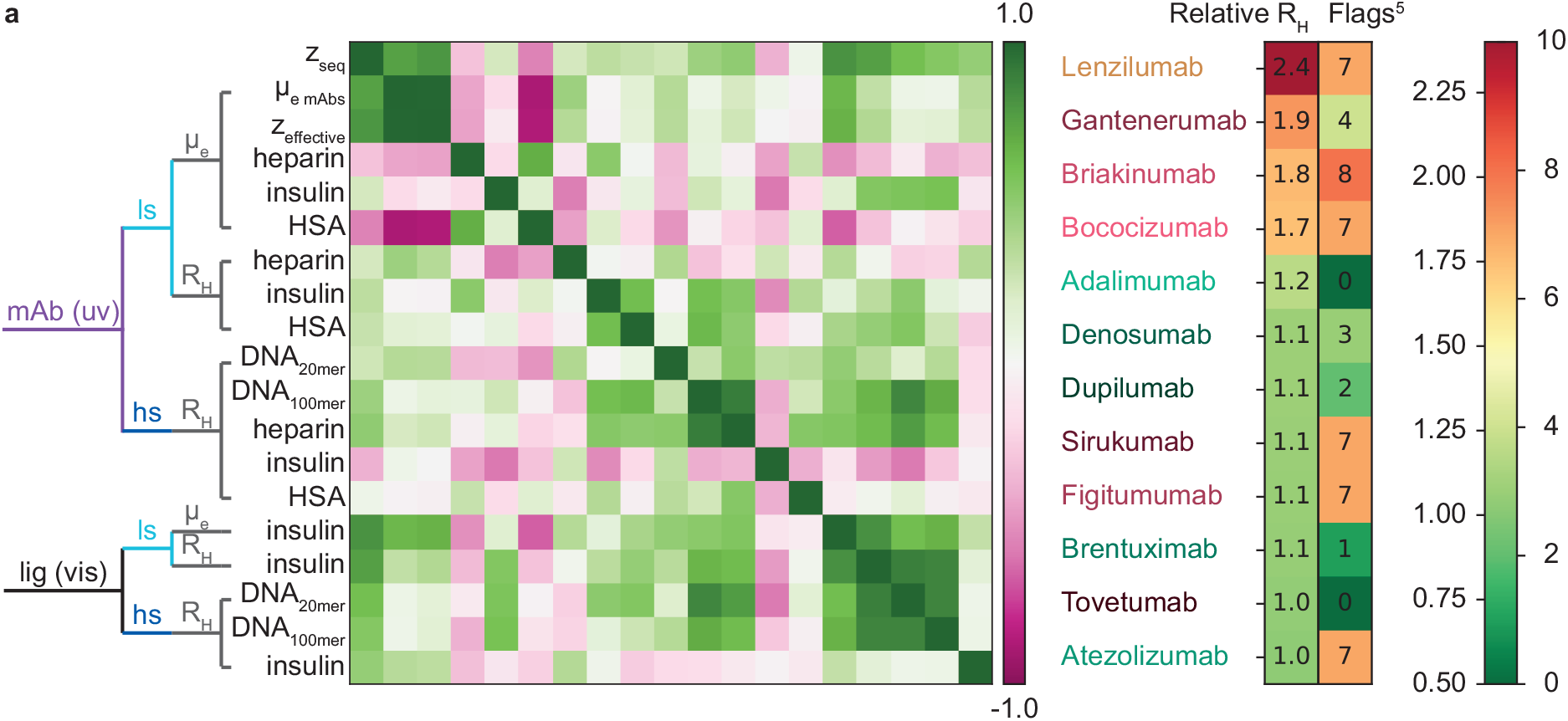
**a** Matrix of Pearson correlation coefficients for the microfluidic screening assays employed in this study. The results are grouped by the observed species (mAb or ligand), the ionic strength (high salt and low salt), and the measured parameter (*R_H_* or *μ_e_*). The physicochemical parameter interrogated for each sample is also shown. **b** Left, ranking of the mAbs by an average of the relative *R_H_* of two non-specificity probes in this study (20mer, hs and insulin, ls), a score of 1 means that no interaction was detected. Right column, number of flags from results in the worst 10% of the panel of 137 mAbs in literature.^5^

#### Improving screening campaigns

Our investigations of the physicochemical properties of antibodies and their non-specific interactions under native solution conditions highlight the need to amend current screening approaches with tools such as the microfluidic platform presented here to obtain accurate and quantitative readouts. We plot the Pearson correlation coefficients for the data collected in this study, Figure 6**a**. The antibody *μ_e_* and *z_e_* are highly correlated due to the similar *R_H_* of the mAbs in this study. The mAb charge correlates with Δ*μ_e_* for binding to heparin and HSA, and insulin interactions with the mAbs, Figure 5.

Monitoring the properties of the non-specificity probe directly increases the sensitivity of the assay, as the mAb *R_H_*/*μ_e_* dominate in most mAb-ligand complexes. A subset of the mAbs (Bo, Br, Ga, Le) emerge as prone to non-specific interactions, Figures 4 and 5. We therefore suggest verifying non-specific binders across two or more targets in solution, e.g. by recording the average relative ligand ratio as a standalone measure of mAb non-specificity, Figure 6**b**. A relative *R_H_* of 1 means that no non-specific binding was detected. Our analysis shows that none of the approved mAbs are flagged with a threshold ≥ 1.2, SI Figure 6**b**. We validate our approach against literature reports for surface- and column-based screens, Figure 2**c-d** and SI Figure 3.^5,23^ Potentially problematic biophysical properties are often flagged as poor performance relative to a cohort, requiring large datasets to define the threshold values, Figure 1**b** and Figure 6**b**.^5^ Quantitative solution-based assays enable us to analyse the non-specificity behaviour of individual antibodies in absolute terms, without using a cohort dataset as benchmark for relative performance. This feature can be particularly useful when assessing the effect of variations in an otherwise similar group of antibodies e.g. variant selection following mutagenesis.

None of the mAbs flagged as problematic in this study have received regulatory approval. Atezolizumab, which is used in cancer treatment, but was flagged as problematic in literature, performed well in the solution-based measurements.^5,21^ Off-target non-specific interactions are one of the potential barriers to mAb developability. This factor is considered together with criteria such as efficacy, cost-benefit, self-association, and specific off-target binding, when deciding whether to advance or discontinue a drug. It is therefore not surprising that three discontinued mAbs gave good results in our non-specificity screens.^37,38^

## Conclusion

Microfluidic solution-based measurements are used to generate a multi-parameter, multicomponent non-specificity fingerprint for a panel of clinical stage antibodies, paving the way for standardised comprehensive physicochemical characterisation of proteins and antibodies. Notably, our approach can identify the physicochemical parameters that drive non-specific interactions, providing a route towards addressing undesirable physicochemical properties before committing to the resource-intensive later stages of development. By recording a quantitative score with a physical meaning based on direct interaction measurements, we find that a subset of the mAbs are prone to non-specific interactions. The solution-based physicochemical measurements in this study provide a well-defined readout and baseline conditions, reducing the risk of false positives compared to surface-based assays such as ELISA. We demonstrate the importance of avidity in enhancing non-specific interactions as an inherent property of the target (e.g. polymers) and experimental setup (e.g. surface immobilisation of the probe).

The solution properties of monoclonal antibodies are key to their success as therapeutics, and their propensity for non-specific interactions is emerging as a central determinant of their developability potential. Quantitative assessments of these parameters provide a valuable addition, not only to the protein drug development pipeline, but also to the wider protein engineering community, and add to our basic knowledge of the physical chemistry governing protein interaction specificity. We envision that the approach outlined here can provide a valuable addition to the state-of-the-art for biophysical chemistry screening workflows.

## Methods

### Microfluidic device preparation

Microfluidic devices were cast in PDMS (Momentive RTV615, Techsil, UK) using standard soft lithography methods^39^. The clear PDMS was coloured black by the addition of a small quantity of carbon nanopowder, 0.2% w/w (Sigma, UK), centrifuged for 40 minutes at 5,000 rcf, air bubbles were removed in a dessicator, and the devices cured at 65 ^*circ*^C for one hour. Inlet and outlet holes were punched using a biopsy punch (WPI, Florida, US). The PDMS devices were bonded to quartz slides (Advalue, US) using a plasma oven to generate the oxygen plasma (Diener Electronics, Germany). The electrodes were fabricated by placing the bonded device microscopy slide down on a hot plate set to 79°C and inserting InBiSn alloy (51%In, 32.5%Bi, 16.5%Sn, Conro Electronics, UK) through the solder inlet.^14^

### Microfluidic diffusional sizing

For each sample, the average diffusion coefficient was determined by imaging the detection region of the diffusional sizing channel to determine the spatial distribution of the sample 1, 16, 40, and 80 mm downstream of its introduction at the centre of the measurement channel (SI Figure 1**a**). Three images were acquired at two different flow rates (40 and 80 *μ*l/hr). The flow rates were set by applying a negative pressure at the outlet using a glass syringe (Hamilton, Switzerland) and controlled by a neMESYS syringe pumps (Cetoni GmbH, Germany). 6 *μ*l sample was loaded for each measurement. Epifluorescence images were acquired with a custom built microscope with a 275 nm LED and a white LED (M275L4, MWWHLP1, and optomechanical parts from Thorlabs, UK), quartz optics (Edmund Optics, UK), a CMOS camera (Prime95B, Photometrics, UK) and Semrock fluorescence filters (UV: TRP-A-000; CF488: ex 475/35, em 525/30, dc 506; Cy3: Cy3-4040C-000, from Laser2000, UK).

The flow rate, device height (individual channels were characterised using a profilometer, Dektak, Brüker), and temperature were taken into account for the data analysis. To determine the spatial sample distribution as a function of time, fluorescence profiles were extracted from the images. As the distance diffused by sample molecules across the channel scales with 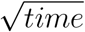, the distances between read points were designed to increase the time interval between read points, thereby enhancing the changes between fluorescence profiles between time points to enable more precise sizing of sample molecules. Diffusion profiles corresponding to the diffusion coefficients of R*_H_* of particles between 0.2 nm and 20 nm were simulated based on laminar Poiseuille flow through the diffusion channel as described previously.^15,40,41^ This range was expanded where the antibodies formed clusters with non-specificity ligands. The best fit to the observed sample distribution at four time points was used to determine the average R*_H_* for each measurement.

### Microfluidic free-flow electrophoresis

Electrophoresis measurements were performed using the custom built microscope described above, which was equipped with a programmable power supply and multimeter as described previously.^14,18^ Buffers and samples were loaded in their respective inlets, and a negative pressure applied at the outlet withdrawal with glass syringe (Hamilton, Switzerland) using neMESYS syringe pumps (Cetoni GmbH, Germany) to control flow rate at 400 *μ*l/hr. As a result of the applied voltage, the sample stream migrated perpendicularly to the direction of flow according to the analyte electrophoretic mobility. The electrophoresis channel used here had a distance between electrodes of 1025 *μ*m and a length of 10,000 *μ*m, the height was measured for each individual channel using a profilometer (Dektak, Brüker). (Figure 5**a**). Three repeats of a voltage range (0 - 1.2 V) with 0.16 V steps were applied for each sample, and at each voltage three images were acquired. Each electrophoretic mobility measurement consumed 8 *μ*L of sample. Independent triplicate measurements of the *μ_e_* were acquired using three different microfluidic chips to analyse each sample composition. The cell constants for individual electrode devices and buffer conductivities were measured using a lock-in amplifier as previously described and a 500 *μ*S cm^−1^ conductivity standard (VWR, UK).^14^ The electrophoresis data was analysed using software written in Python.^17^ Linear fits to the slope of the plots of velocity against electric field, which were used to determine *μ_e_* for each sample.

### Sample preparation

Standard buffer components were purchased from Sigma. Buffer solutions were degassed prior to use to prevent the introduction of bubbles into the microfluidic channels. HSA and heparin were purchased (A3782 Sigma Aldrich and A3004,001 from Applichem). DNA oligomers were purchased with a 5’ Cy3 modification (100, 20 mer: Merck, Germany, HPLC purified and lyophilised; 10, 6 mer: Biomers, Germany, HPLC purified and lyophilised). The sequences used were: 100mer - CTCACCCACAACCACAAACAATTTAAATAATATTAAATAATATTAATATATTATCGATTAAATAATAATTAATTAATATTGGTTGGATGGTAGATGGTGA; 20mer - CTCACCCACAACCACAAACA; 10mer - TATACTTAGT; 6mer - ATGCGA. Antibodies and monomeric insulin were prepared as outlined below.

### Antibody expression and purification

The primary sequence of the clinical-stage antibodies in IgG1 subclass format where taken from the paper by Jain et al.^5^ Expression plasmids for production were purchased from Twist Bioscience. All compounds were expressed by transient transfection of Expi293 cells following the instructions of the manufacturer (Expi system, Life technologies) and the resulting compounds were purified from cell culture supernatants using an kta Express chromatography system using affinity purification using Mabselect Sure Protein A resin followed by a gel filtration column based on the Superdex200 resin (GE Healthcare). The MabSelect Sure Protein A column was washed with 0.1 M Hepes, 150 mM NaCl, pH 7.4 and antibodies were eluted from the affinity column with 0.1 M sodium formate, pH 3.5 onto the pre-equilibrated gel filtration column via an interconnected loop. The gel filtration column was operated using a running buffer based on 20 mM HEPES, 0.15 M NaCl, pH 7.4. Eluted peaks were fractionated in 96-well plates and fractions were pooled to obtain high purities with minimum of high-molecular weight protein.

### Labelling of monomeric insulin

2mM of recombinantly produced and purified human insulin (B28D) was incubated with 2.5 mM of CF488A succinimidyl ester in a phosphate buffer at pH 8 for 4 hours at RT. Reaction was stopped with Tris and sample purified on a gel filtration column to get rid of excess probe. LC-MS analysis showed both unlabelled and 1:1 probe labelled insulin. Spectrophotometric quantification of the sample resulted in a Degree of Labelling of approximately 50%.

## Supporting information

Supplementary Information: Non-specificity fingerprints for clinical stage antibodies in solution

## Acknowledgements

We would like to thank Dr Tushar Jain and Prof. Dane Wittrup (Adimab and MIT) for sharing the individual ELISA readouts for DNA and insulin ligands and allowing us to show them together with our solution-based measurements in this paper. We also thank Dr Laila Sakhnini (Novo Nordisk A/S) for great discussions during the project. In addition to the collaboration between Novo Nordisk A/S and the University of Cambridge, we would like to acknowledge financial support from the Newman Foundation (T.P.J.K.).

## Data

Data available at https://doi.org/10.6084/m9.figshare.21858645.

