## Supplementary Information: Non-specificity fingerprints for clinical stage antibodies in solution for "Non-specificity fingerprints for clinical stage antibodies in solution"

A dozen monoclonal antibodies (mAbs) were selected for this study based on their reported biophysical properties, sequence charge, and approval status.

Five of the mAbs have received regulatory approval:

- **Adalimumab** targets tumour necrosis factor $\alpha$ (TNF$\alpha$). Adalimumab is approved as a subcutaneous injection for the treatment of a number of autoimmune conditions including rheumatoid arthritis, Chron's disease, plaque psoriasis, and ulcerative cholitis.[1] It has a good biophysical profile, and was not flagged in the screen by Jain et al.[2] We included Adalimumab in this study as a well-behaved control to compare with the mAbs that have a poor biophysical profile in literature.
- **Atezolizumab** targets programmed death-ligand 1 (PD-L1) and has been approved in can-

cer treatments including non-small cell lung cancer, small cell lung cancer, breast cancer, hepatocellular carcinoma, and urothelial carcinoma.[1] It is administered as an intravenous infusion. Interestingly, Atezolizumab performed poorly across the biophysical assays employed by Jain et al. (Figure 1).[2] We were therefore interested to see whether this poor behaviour translated to solution assays.

- **Brentuximab** targets CD30 and is approved as an intravenous infusion for the treatment of CD30 positive Hodgkin lymphoma, systemic anaplastic large cell lymphoma, and CD30 positive T-cell lymphoma.[1,3] We chose to investigate this mAb as it had been flagged in surface-based assays, but otherwise performed well in literature.[2] Brentuximab would therefore be an interesting case for a comparison of surface and solution-based non-specificity screens. In addition, the sequence charge of Brentuximab is near neutral.

- **Denosumab** targets receptor activator of nuclear factor-$\kappa$B ligand (RANKL). It is approved as a subcutaneous injection in the treatment of osteoperosis and for the prevention of skeletal related events in patients with bone metastasis.[1,4] Denosumab was flagged in a number of surface-based assays and we therefore included it in our study.[2]

- **Dupilumab** targets interleukin 4 (IL-4) and interleukin 13 (IL-13). It is approved as a subcutaneous injection for the treatment of a number of autoimmune conditions including atopic eczema, severe asthma with type 2 inflammation, and severe chronic rhinosinusitis with nasal polyps.[1] Dupilumab was flagged in surface-based assays only by Jain et al. and therefore included in this study.[2]

One of the mAbs is currently in clinical trials:

- **Lenzilumab** targets granulocyte macrophage colony stimulating factor (GMC SF) and is currently being investigated for the treatment of COVID-19, clinicaltrials.gov identifier NCT04351152.[5] Lenzilumab is the mAb in this study with the most +ve sequence charge.[2]

Six of the mAbs have been discontinued:

- **Bococizumab** targets proprotein convertase subtilisin/kexin type 9 (PCSK9) and was discontinued following clinical trials.[6] The mAb was included in this study due to poor biophysical properties in literature.[2]
- **Briakinumab** targets interleukins 12 and 23 (IL-12, IL-23) and was withdrawn after clinical trials.[7] We included Briakinumab here in the group of mAbs with poor biophysical properties from previous reports.[2]
- **Figitumumab** targets insulin-like growth factor 1 receptor (IGF-1R). Development of Figitumumab was discontinued after patients had shorter overall survival in the Figitumumab group in clinical trials.[8] The mAb was included here in the poor reported biophysical properties and outcomes group.[2]
- **Gantenerumab** targets amyloid-$\beta$42 fibrils (A$\beta$42 fibrils) and has recently been discontinued after clinical trials.[9–11] The mAb was flagged in some assays by Jain et al. but is not amongst the cohort with the worst biophysical properties. We included it here as the

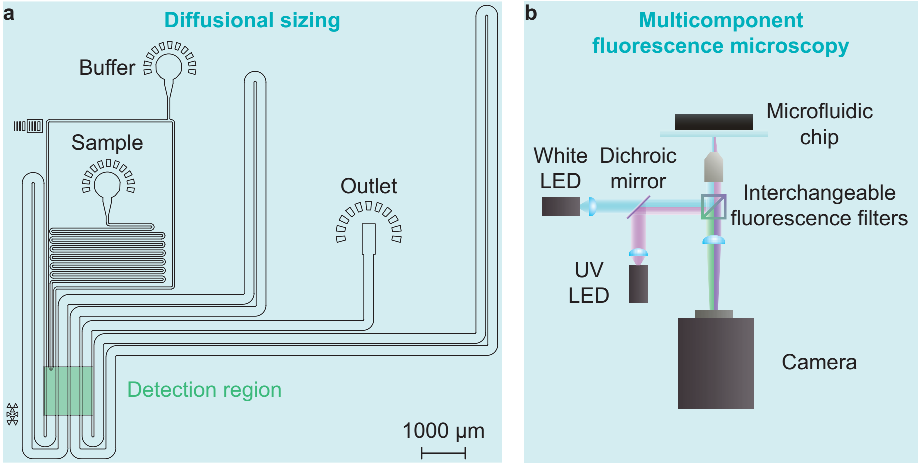


**Figure 1:** Precipitation of Adalimumab in the presence of DNA oligomers of different lengths in ls buffer easily forms a pellet, even when spun for a minute in a bench top mini centrifuge (from the left: 20, 50, 100 nucleotides).

non-fibrillar A$\beta$42 peptide is intrinsically disordered, and A$\beta$42 could therefore represent a particularly challenging target with respect to non-specificity. Additionally, Gantenerumab was included here due to its highly positive sequence charge.

- **Sirukumab** targets interleukin 6 (IL-6) and has recently investigated for the treatment of COVID-19.[12,13] Sirukumab was not approved by the FDA following previous clinical trials for the treatment of rheumatoid arthritis[14,15]

- **Tovetumab** targets platelet-derived growth factor receptor $\alpha$ (PDGFR) and did not meet its primary endpoint in clinical trials.[16] Recent studies have focussed on elucidating the mechanism and pharmacokinetics of Tovetumab.[17] We included Tovetumab in this study because it has good reported biophysical properties without being successful in clinical trials.[2,16]

| Antibody | Antigen | $R_H$ hs / nm | $R_H$ ls / nm | $\mu_e$ ls / $10^{-8}$ m V$^{-1}$ s$^{-1}$ | $z_e$ ls / e |
|---|---|---|---|---|---|
| Ad, Adalimumab | TNF$\alpha$ | 5.6 $\pm$ 0.14 | 5.6 $\pm$ 0.56 | 0.61 $\pm$ 0.05 | 3.8 $\pm$ 0.49 |
| Az, Atezolizumab | PD-L1 | 6.0 $\pm$ 0.25 | 5.7 $\pm$ 0.11 | 0.37 $\pm$ 0.06 | 2.4 $\pm$ 0.37 |
| Bo, Bococizumab | PCSK9 | 6.4 $\pm$ 0.48 | 5.8 $\pm$ 0.41 | 0.86 $\pm$ 0.16 | 5.4 $\pm$ 1.2 |
| Br, Briakinumab | IL-12, IL-23 | 5.8 $\pm$ 0.28 | 5.9 $\pm$ 0.48 | 0.53 $\pm$ 0.12 | 3.5 $\pm$ 0.85 |
| Bx, Brentuximab | CD30 | 6.2 $\pm$ 0.54 | 5.8 $\pm$ 0.38 | -0.01 $\pm$ 0.06 | -0.09 $\pm$ 0.29 |
| De, Denosumab | RANKL | 5.8 $\pm$ 0.41 | 5.2 $\pm$ 0.22 | 0.74 $\pm$ 0.07 | 4.3 $\pm$ 0.37 |
| Du, Dupilumab | IL-4, IL-13 | 6.0 $\pm$ 0.33 | 5.4 $\pm$ 0.19 | 0.57 $\pm$ 0.04 | 3.5 $\pm$ 0.22 |
| Fi, Figitumumab | IGF-1R | 5.8 $\pm$ 0.32 | 6.0 $\pm$ 0.42 | 0.63 $\pm$ 0.10 | 4.2 $\pm$ 0.64 |
| Ga, Gantenerumab | A$\beta$42 | 6.0 $\pm$ 0.21 | 5.9 $\pm$ 0.47 | 0.70 $\pm$ 0.16 | 4.7 $\pm$ 0.96 |
| Le, Lenzilumab | GM-CSF | 6.4 $\pm$ 0.43 | 6.5 $\pm$ 0.59 | 0.51 $\pm$ 0.20 | 3.7 $\pm$ 1.2 |
| Si, Sirukumab | IL-6 | 6.2 $\pm$ 0.39 | 5.8 $\pm$ 0.14 | 0.31 $\pm$ 0.03 | 2.0 $\pm$ 0.18 |
| To, Tovetumab | PDGFR$\alpha$ | 6.3 $\pm$ 0.35 | 6.1 $\pm$ 0.64 | 0.75 $\pm$ 0.06 | 5.1 $\pm$ 0.62 |

**Table 1:** Table of the measured solution state properties (1 mg/ml mAbs).

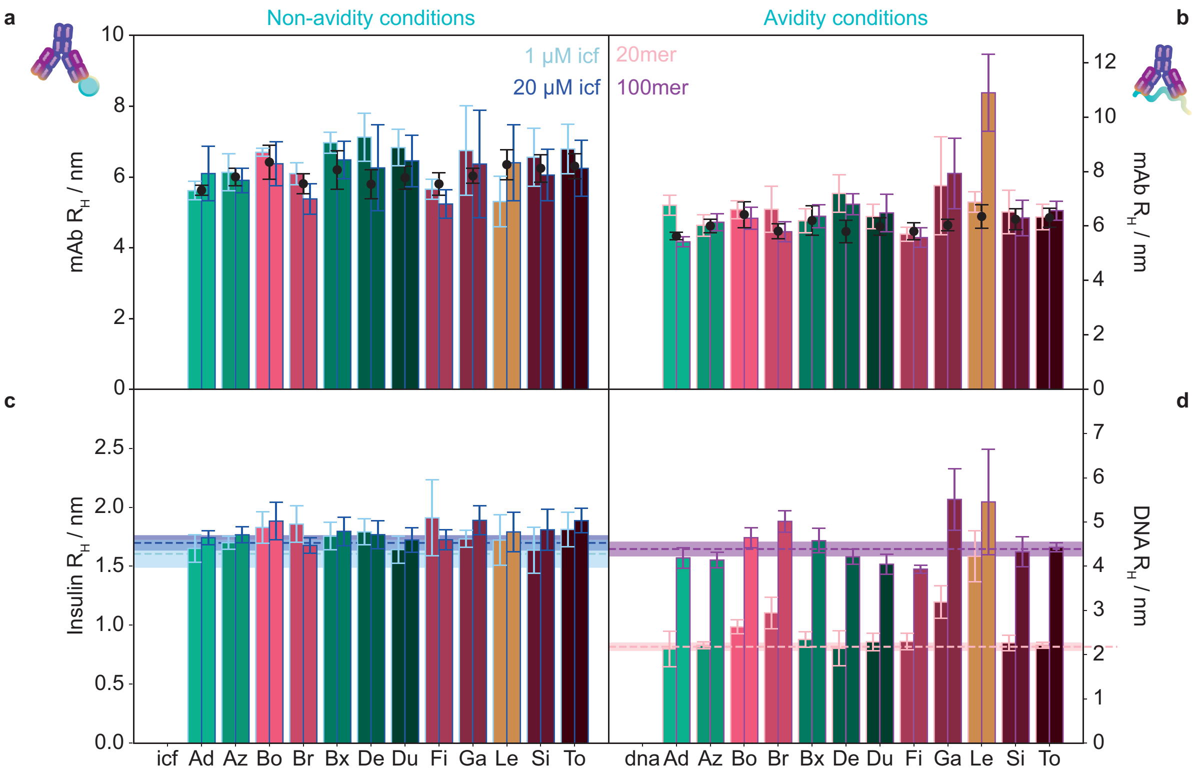


**Figure 2:** **a** Intrinsic fluorescence data: $R_H$ for non-specific binding assayed in free solution under non-avidity conditions in the high salt buffer using 1 $\mu$M and 20 $\mu$M insulin-CF488 and 6.7 $\mu$M mAb. The antibody alone is shown in black. **b** Intrinsic fluorescence data: Interactions where target avidity is possible with single-stranded DNA polymers (20mer and 100mer, Cy3-labelled). **c** Ligand fluorophore fluorescence data: non-specific binding assayed in free solution under non-avidity conditions in the high salt buffer using 1 $\mu$M and 20 $\mu$M insulin-CF488 and 6.7 $\mu$M mAb. **d** Ligand fluorophore fluorescence data: Interactions where target avidity is possible with single-stranded DNA polymers (20mer and 100mer, Cy3-labelled).

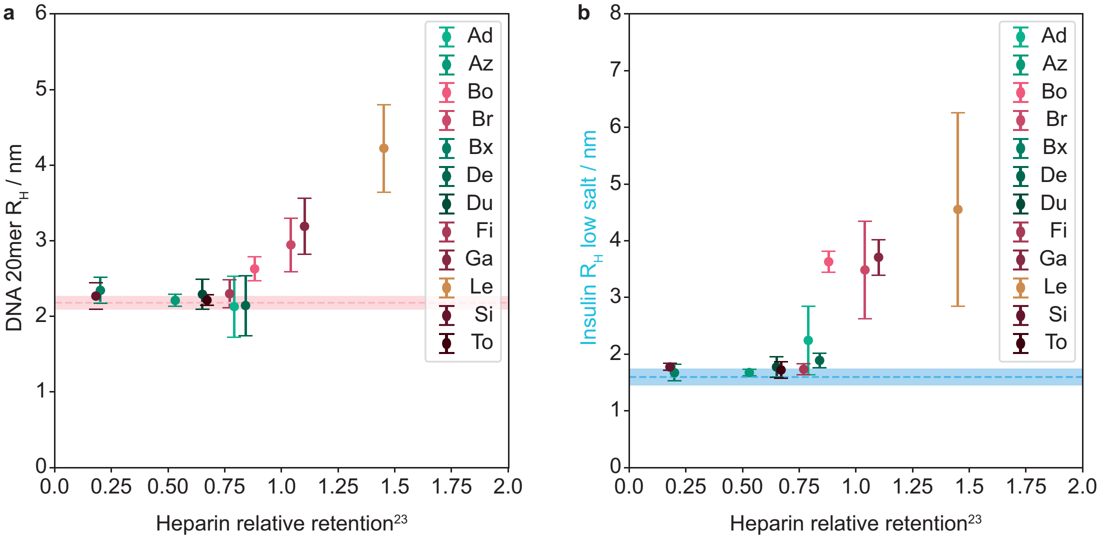


**Figure 3:** **a** Ligand $R_H$ for DNA (1 $\mu$M 20mer, Cy3-labelled) measured in the presence of 1 mg/ml mAb in high salt buffer against the relative heparin retention reported by Kraft et al. for the mAb panel (reference 23 in the main text).[18] **b** Ligand $R_H$ for monomeric insulin (1 $\mu$M, CF488-labelled) measured in the presence of 1 mg/ml mAb in low salt buffer against the relative heparin retention reported by Kraft et al. in their column-based assay.[18]


1. Dingfelder, F., Henriksen, A., Wahlund, P.-O., Arosio, P. & Lorenzen, N. Measuring Self-AssociationSelf-association of AntibodyAntibodiesLead Candidates with Dynamic Light ScatteringDynamic light scattering (DLS). In Houen, G. (ed.) *Therapeutic Antibodies: Methods and Protocols*, 241–258 (Springer US, New York, NY, 2021).

2. Jain, T. *et al.* Biophysical properties of the clinical-stage antibody landscape. *Proceedings of the National Academy of Sciences* **114**, 944–949 (2017).

3. Weyden, C. A. V. D., Pileri, S. A., Feldman, A. L., Whisstock, J. & Prince, H. M. Understanding CD30 biology and therapeutic targeting : a historical perspective providing insight into future directions. *Nature Publishing Group* 1–10 (2017).

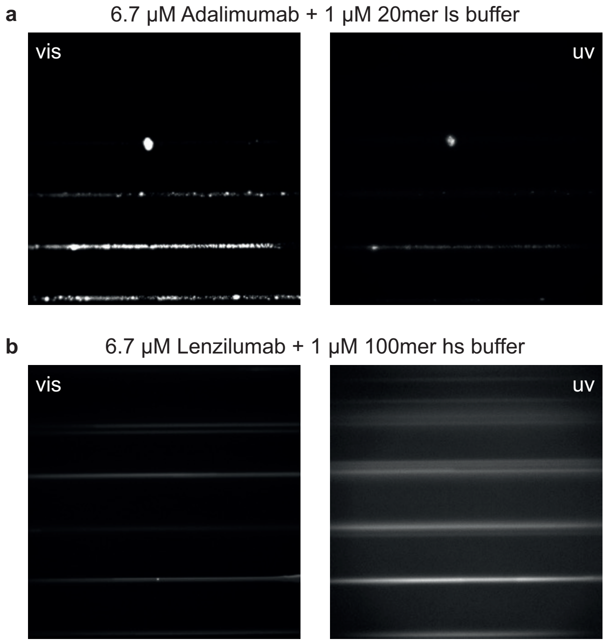


**Figure 4:** **a** Fluorescence images of cluster formation between Adalimumab and ssDNA 20mer under low salt conditions. Visible fluorescence imaging (vis, left) of the labelled DNA oligomer and intrinsic fluorescence from the antibody (uv, right) show that both species are incoporated in the clusters, and that these particles are deposited in the microfluidic channel. **b** Fluorescence images of cluster formation between Lenzilumab and ssDNA 100mer under high salt conditions. Visible fluorescence imaging (vis, left) of the labelled DNA oligomer and intrinsic fluorescence from the antibody (uv, right) show that both species are incoporated in the clusters.

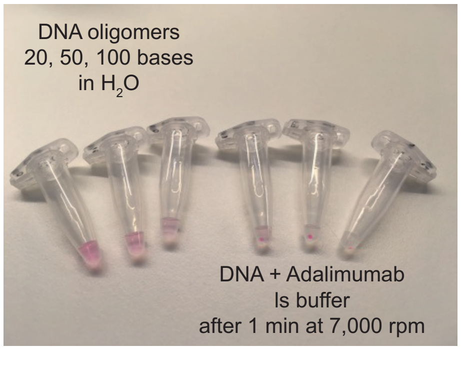


**Figure 5:** Precipitation of Adalimumab in the presence of DNA oligomers of different lengths in ls buffer easily forms a pellet, even when spun for a minute in a bench top mini centrifuge (from the left: 20, 50, 100 nucleotides).

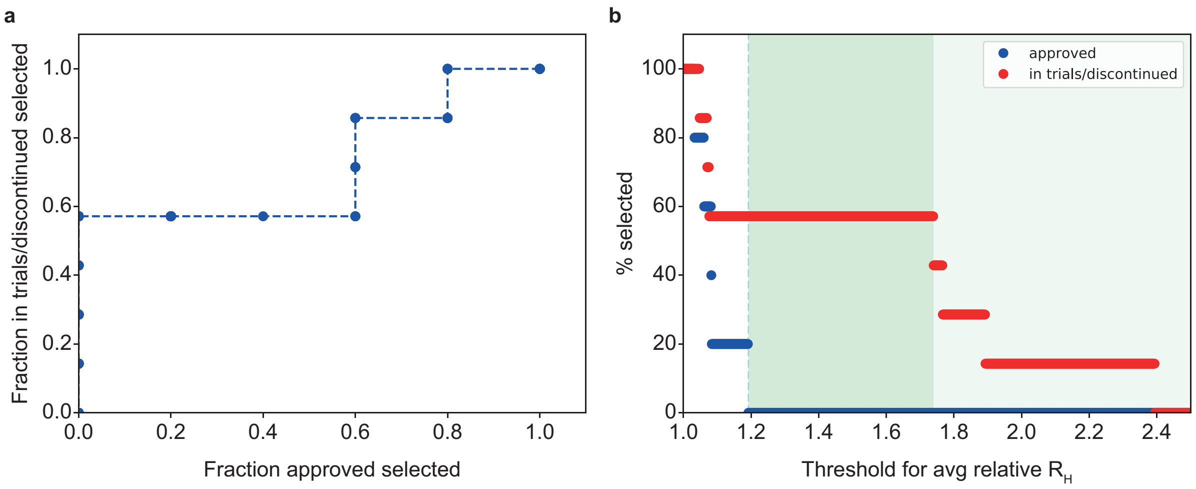


**Figure 6: a** Receiver operator characteristic (ROC) curve when using the average relative $R_H$ to classify mAb non-specificity ($R_H$ for 20mer in high salt and insulin in low salt buffer). The approved mAbs are classed as false positives. Here we show the fraction of false positives for non-specificity (approved mAbs) against the fraction of potentially true positives (discontinued mAbs and those in clinical trials). **b** The percentage of approved and discontinued/in trials mAbs selected against threshold value for the average rel $R_H$ for two non-specificity targets (20mer in high salt and insulin in low salt buffer). Threshold values where no approved mAbs are flagged are shaded in green.

11. Travis, J. Roche Alzheimer's antibody fails to slow cognitive decline in major test. *Science* (2022).
